## Supplementary Information for "Collagen breaks at weak sacrificial bonds taming its mechanoradicals"

October 17, 2022

### 1 Supplementary figures and data tables

#### 1.1 Data for the PYD crosslink

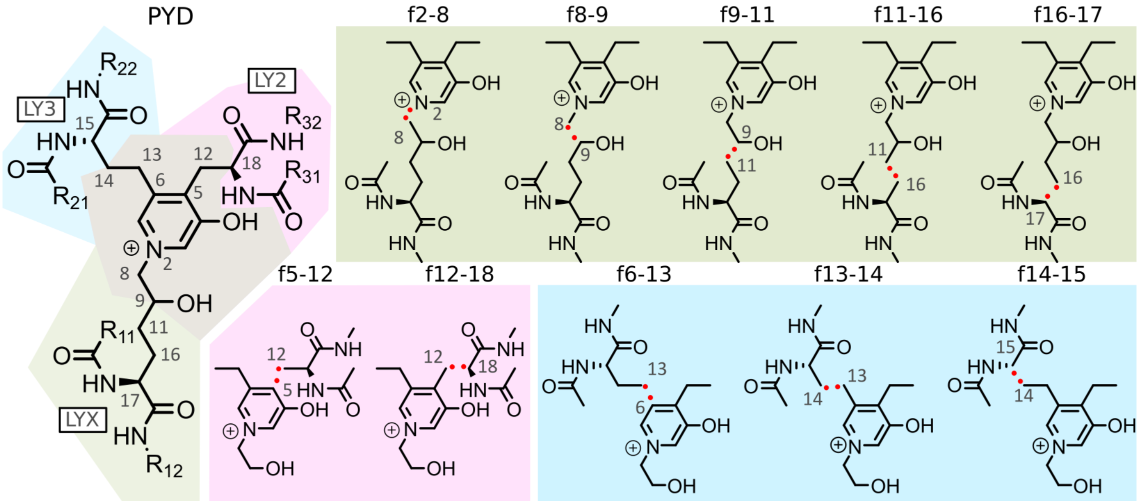


Suppl. Fig. 1: **Internal nomenclature of molecular structures used for BDE calculations of the PYD crosslink.** The presented nomenclature corresponds to the following Suppl. Tables 1-2. The "a" fragment always includes the central nitrogen, the "b" fragment always the truncated backbone.

| Molecule | ZPVE [Ha] | H0K [Ha] | H298K [Ha] |
|---|---|---|---|
| 0_PYD_b | -1167.81640 | -1167.34701 | -1167.31685 |
| 1_PYD_l | -1089.26818 | -1088.85407 | -1088.82669 |
| 2_PYD_r | -1049.99417 | -1049.60825 | -1049.58210 |
| f11-16b | -494.71218 | -494.54312 | -494.52969 |
| f11-16a | -672.95343 | -672.66416 | -672.64603 |
| f14-15a | -633.67634 | -633.41467 | -633.39786 |
| f5-12a | -555.09102 | -554.88329 | -554.86945 |
| f13-14a | -594.41633 | -594.18073 | -594.16598 |
| f12-18a | -594.41509 | -594.17963 | -594.16479 |
| f12-18a depr | -594.02805 | -593.80541 | -593.79083 |
| f2-8b | -687.69188 | -687.43480 | -687.41585 |
| f2-8a | -479.94870 | -479.74752 | -479.73474 |
| f6-13a | -555.10169 | -554.89306 | -554.87969 |
| f16-17b | -455.46470 | -455.32179 | -455.30981 |
| f16-17a | -712.22699 | -711.90990 | -711.89039 |
| f9-11a | -633.68808 | -633.42542 | -633.40879 |
| f9-11b | -533.98481 | -533.78833 | -533.77329 |
| f8-9b | -648.42901 | -648.19758 | -648.18042 |
| f8-9a | -519.23885 | -519.00942 | -518.99564 |
| f8-9a-H-capped | -519.90996 | -519.66678 | -519.65253 |
| f8-9b-H-capped | -649.09141 | -648.84678 | -648.82959 |
| f12-18-depr-a-H-capped | -594.66872 | -594.43435 | -594.41912 |
| f16-17a-H-capped | -712.89938 | -712.56813 | -712.54870 |
| f16-17b-H-capped | -456.11102 | -455.95560 | -455.94338 |
| f11-16b-H-capped | -495.38826 | -495.20462 | -495.19139 |
| f11-16a-H-capped | -673.62836 | -673.32479 | -673.30686 |
| f6-13a-H-capped | -555.80212 | -555.58054 | -555.56716 |
| f14-15a-H-capped | -634.35094 | -634.07416 | -634.05775 |
| f9-11a-H-capped | -634.35030 | -634.07402 | -634.05750 |
| f9-11b-H-capped | -534.65818 | -534.44669 | -534.43203 |
| f5-12a-H-capped | -555.79213 | -555.57170 | -555.55788 |
| f13-14a-H-capped | -595.07984 | -594.83101 | -594.81599 |
| f2-8a-H-capped | -480.63884 | -480.42239 | -480.40999 |
| f2-8b-H-capped | -688.36708 | -688.09511 | -688.07657 |
| f12-18a-H-capped | -595.07251 | -594.82470 | -594.80933 |

Suppl. Table 1: **G4(MP2)-6X energies of radicals, reference structures and fragments of the PYD crosslink.** The nomenclature follows Suppl. Fig. 1.

| Bond | BDE | Rad. (a) | center (a) | RSE (a) | Rad. (b) | center (b) | RSE (b) | RSE sum |
|---|---|---|---|---|---|---|---|---|
| f12-18 | 282.2 | f12-18a | C | -58.3 | f12-18b | C | -87.1 | -145.4 |
| f12-18 depr | 220.0 | f12-18a depr | C | -101.0 | f12-18b | C | -87.1 | -188.1 |
| f16-17 | 306.3 | f16-17a | C | -22.2 | f16-17b | C | -87.1 | -109.3 |
| f14-15 | 312.5 | f14-15a | C | -18.0 | f14-15b | C | -87.1 | -105.1 |
| f13-14 | 344.0 | f13-14a | C | -43.9 | f13-14b | C | -13.2 | -57.1 |
| f9-11 | 353.8 | f9-11a | C | -47.4 | f9-11b | C | -21.0 | -68.4 |
| f8-9 | 369.7 | f8-9a | C | -25.9 | f8-9b | C | -46.1 | -72.0 |
| f11-16 | 370.5 | f11-16a | C | -15.5 | f11-16b | C | -13.2 | -28.7 |
| f2-8 | 436.5 | f2-8a | N | 13.7 | f2-8b | C | -15.8 | -2.1 |
| f6-13 | 456.1 | f6-13a | C | 54.4 | f6-13b | C | -21.0 | 33.4 |
| f5-12 | 480.3 | f5-12a | C | 56.9 | f5-12b | C | -13.2 | 43.7 |

Suppl. Table 2: **BDEs, RSEs, and summed RSEs of PYD bonds and radicals.** The nomenclature follows Suppl. Fig. 1. All values are given in kJ/mol. BDEs were calculated from enthalpies at 298 K (H298K).

#### 1.2 Data for DPD crosslink

The absent hydroxyl group in DPD did not affect the $C_\alpha$-$C_\beta$ bond of the top right arm, implying the same for the $C_\alpha$-$C_\beta$ bond of the top left arm. The $C_\alpha$-$C_\beta$ bond of the lower arm is closer to the hydroxyl group, but the effect on bonds in $\gamma$ position should be on the order of the methodological error. Hence, we do not differentiate between low BDEs of PYD and DPD in the manuscript.

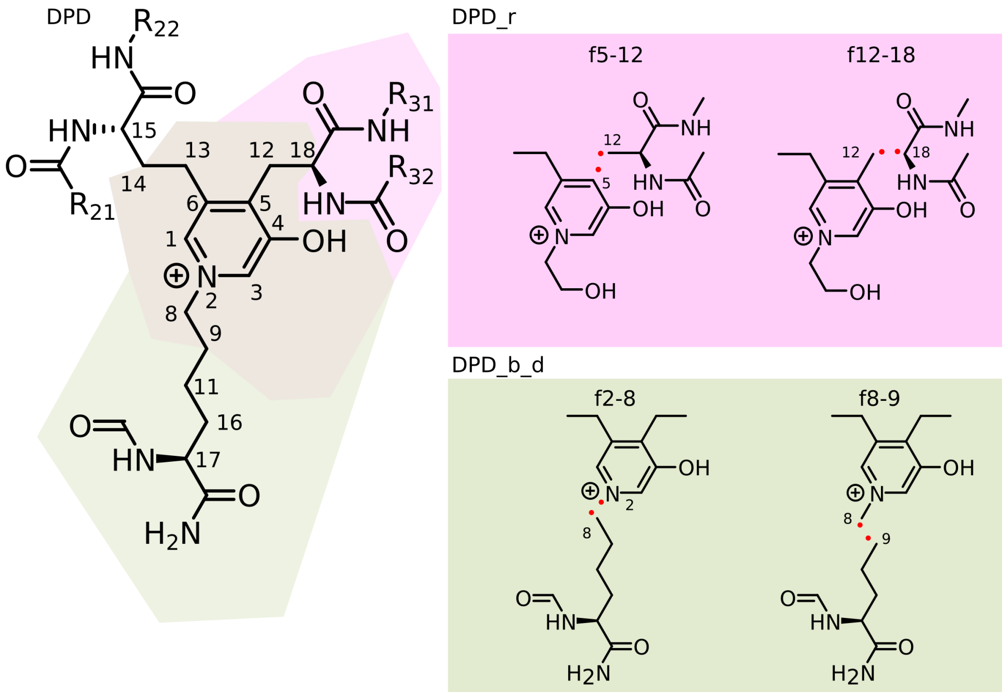


Suppl. Fig. 2: **Internal nomenclature of molecular structures used for calculations of the DPD crosslink.** The presented nomenclature corresponds to the following Suppl. Tables 3 and 4. The "a" fragment always includes the central nitrogen, the "b" fragment always the truncated backbone.

| Molecule | ZPVE [Ha] | H0K [Ha] | H298K [Ha] |
|---|---|---|---|
| 1_DPD_b_d | -1014.10090 | -1013.69091 | -1013.66534 |
| 2_DPD_r | -974.83586 | -974.45517 | -974.43024 |
| f2-8b | -533.98242 | -533.78462 | -533.77025 |
| f8-9b | -494.71165 | -494.54175 | -494.52878 |
| f5-12a | -479.93279 | -479.73017 | -479.71751 |
| f12-18a | -519.25667 | -519.02679 | -519.01303 |
| f2-8a | -479.94870 | -479.74752 | -479.73474 |
| f8-9a | -519.23885 | -519.00942 | -518.99564 |
| f11-16b | -494.71218 | -494.54312 | -494.52969 |
| f16-17b | -455.46470 | -455.32179 | -455.30981 |

Suppl. Table 3: **G4(MP2)-6X energies of radicals, reference structures and fragments of the DPD crosslink.** The nomenclature follows Suppl. Fig. 2.

| Bond | BDE | Rad. (a) | center (a) | Rad. (b) | center (b) |
|---|---|---|---|---|---|
| f2-8 | -421.0 | f2-8a | N | f2-8b | C |
| f8-9 | -370.0 | f8-9a | C | f8-9b | C |
| f5-12 | -480.6 | f5-12a | C | f5-12b | C |
| f12-18 | -282.0 | f12-18a | C | f12-18b | C |

Suppl. Table 4: **BDEs of DPD bonds.** The nomenclature follows Suppl. Fig. 2. All values are given in kJ/mol. BDEs were calculated from enthalpies at 298 K (H298K).

#### 1.3 Data for HLKNL crosslink

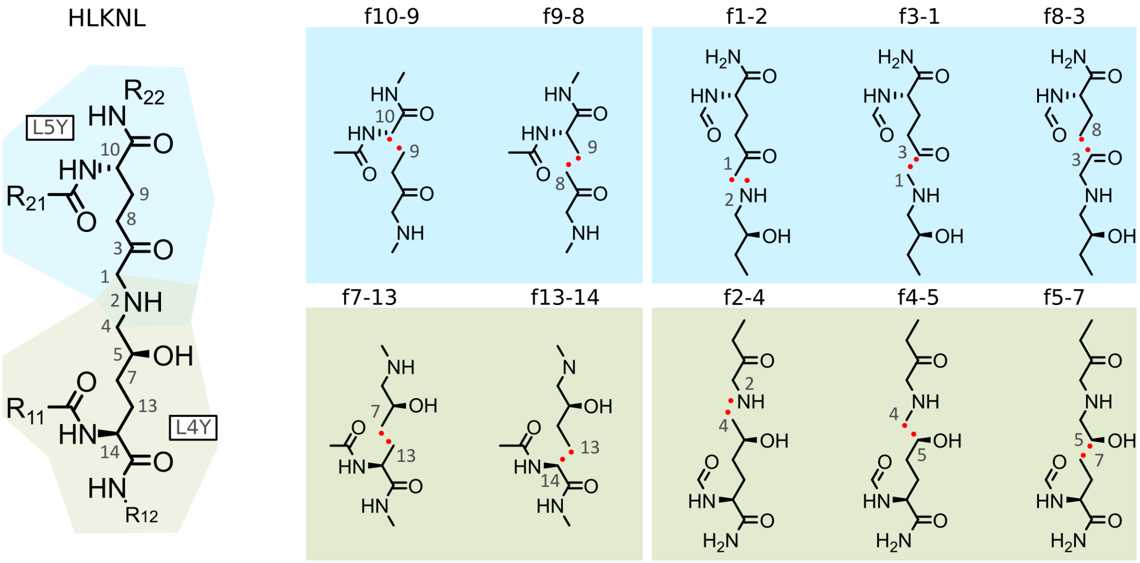


Suppl. Fig. 3: **Internal nomenclature of molecular structures used for BDE calculations of the HLKNL crosslink.** The presented nomenclature corresponds to the Suppl. Tables 5 and 6. The "a" fragment always includes the central nitrogen, the "b" fragment always the truncated backbone.

| Molecule | ZPVE [Ha] | H0K [Ha] | H298K [Ha] |
|---|---|---|---|
| p_hlknl_t_long_prod | -896.14927 | -895.82246 | -895.79977 |
| p_hlknl_b_long_prod | -896.15996 | -895.83177 | -895.80962 |
| p_hlknl_t_short_prod | -781.71295 | -781.41905 | -781.39826 |
| p_hlknl_b_short_prod | -782.92794 | -782.61089 | -782.58965 |
| f8-9b | -569.88602 | -569.70896 | -569.69538 |
| f1-3a | -327.35260 | -327.18660 | -327.17616 |
| f1-3b | -568.67824 | -568.52396 | -568.51081 |
| fd9-11b | -455.44184 | -455.29971 | -455.28825 |
| f1-2a | -288.07841 | -287.94104 | -287.93204 |
| f2-8b | -609.14869 | -608.94587 | -608.93055 |
| f4-5a | -326.13899 | -325.99668 | -325.98663 |
| f5-7a | -440.57469 | -440.39845 | -440.38636 |
| f2-4a | -286.86604 | -286.75235 | -286.74373 |
| f1-2b | -607.94753 | -607.76695 | -607.75251 |
| f3-8a | -440.57027 | -440.39406 | -440.38162 |
| f10-9a | -326.12088 | -325.97975 | -325.96937 |
| f8-9a | -286.86199 | -286.74703 | -286.73867 |
| f13-14a | -327.33663 | -327.17178 | -327.16119 |
| f7-13a | -288.06311 | -287.92611 | -287.91678 |
| f11-16b | -494.71218 | -494.54312 | -494.52969 |
| f16-17b | -455.46470 | -455.32179 | -455.30981 |
| f16-17b-H-capped | -456.11102 | -455.95560 | -455.94338 |
| f2-4b-H-capped | -609.82541 | -609.60794 | -609.59294 |
| f5-7b-H-capped | -456.11584 | -455.95882 | -455.94769 |
| f5-7a-H-capped | -441.23646 | -441.04690 | -441.03479 |
| f11-16b-H-capped | -495.38826 | -495.20462 | -495.19139 |
| f10-9a-H-capped | -326.79839 | -326.64264 | -326.63267 |
| f2-4a-H-capped | -287.53826 | -287.40996 | -287.40128 |
| f13-14a-H-capped | -328.00952 | -327.83033 | -327.81987 |
| f1-2a-H-capped | -288.75171 | -288.59974 | -288.59068 |
| f4-5a-H-capped | -326.79839 | -326.64264 | -326.63267 |
| f3-8a-H-capped | -441.22368 | -441.03554 | -441.02313 |
| f8-9a-H-capped | -287.52586 | -287.39808 | -287.38935 |
| f1-3b-H-capped | -569.33021 | -569.16414 | -569.15097 |
| f4-5b-H-capped | -570.54905 | -570.35865 | -570.34509 |
| f1-3a-H-capped | -328.00658 | -327.82769 | -327.81712 |
| f7-13a-H-capped | -288.73962 | -288.58827 | -288.57920 |
| f1-2b-H-capped | -608.61124 | -608.41755 | -608.40282 |

Suppl. Table 5: **G4(MP2)-6X energies of radicals, reference structures and fragments of HLKNL crosslink.** The nomenclature follows Suppl. Fig. 3.

| Bond | BDE | Rad. (a) | center (a) | RSE (a) | Rad. (b) | center (b) | RSE (b) | RSE sum |
|---|---|---|---|---|---|---|---|---|
| f1-3 | 296.1 | f1-3a | C | -67.7 | f1-3b | C | -69.8 | -137.5 |
| f1-2 | 302.5 | f1-2a | N | -29.9 | f1-2b | C | -43.2 | -73.1 |
| f13-14 | 311.5 | f13-14a | C | -21.2 | f13-14b | C | -87.1 | -108.3 |
| f10-9 | 312.6 | f10-9a | C | -9.1 | f10-9b | C | -87.1 | -96.2 |
| f4-5 | 335.0 | f4-5a | C | -54.4 | f4-5b | C | -44.7 | -99.1 |
| f8-9 | 341.0 | f8-9a | C | -42.2 | f8-9b | C | -13.2 | -55.4 |
| f3-8 | 341.1 | f3-8a | C | -66.2 | f3-8b | C | -19.2 | -85.4 |
| f5-7 | 354.5 | f5-7a | C | -48.1 | f5-7b | C | -19.2 | -67.3 |
| f2-4 | 355.3 | f2-4a | N | -32.8 | f2-4b | C | -11.4 | -44.2 |
| f7-13 | 375.9 | f7-13a | C | -11.4 | f7-13b | C | -13.2 | -24.6 |

Suppl. Table 6: **BDEs, RSEs, and summed RSEs of HLKNL bonds and radicals.** The nomenclature follows Suppl. Fig. 3. All values are given in kJ/mol. BDEs were calculated from enthalpies at 298 K (H298K).

#### 1.4 Data for deH-LNL bonds

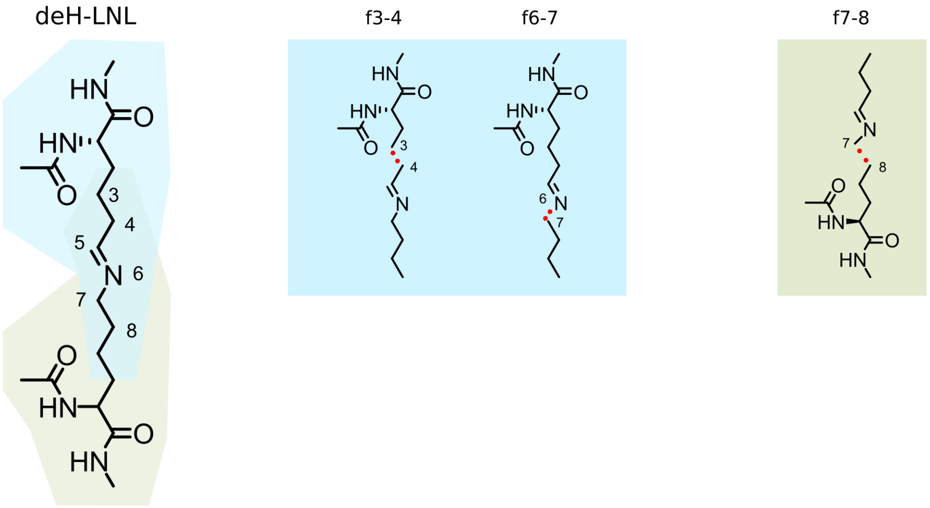


Suppl. Fig. 4: **Internal nomenclature of molecular structures used for BDE calculations of the deH-LNL crosslink.** The presented nomenclature corresponds to the Suppl. Tables 7 and 8. The "a" fragment always includes the central nitrogen, the "b" fragment always the truncated backbone.

| Molecule | ZPVE [Ha] | H0K [Ha] | H298K [Ha] |
|---|---|---|---|
| p_deHLNL_LX5_long_prod | -745.80936 | -745.49182 | -745.47156 |
| p_deHLNL_LX4_long_prod | -745.80937 | -745.49200 | -745.47170 |
| f6-7a-LX4 | -211.70237 | -211.59250 | -211.58504 |
| f6-7b-LX4 | -533.98211 | -533.78406 | -533.76966 |
| f7-8a | -250.96554 | -250.82925 | -250.82035 |
| f6-7a-LX5 | -588.07866 | -587.88532 | -587.87073 |
| f7-8b | -494.71125 | -494.54110 | -494.52803 |
| f6-7b-LX5 | -157.60522 | -157.49077 | -157.48347 |
| f3-4a | -290.23638 | -290.07138 | -290.06134 |
| f3-4b | -455.44184 | -455.29971 | -455.28825 |
| f6-7b-H-capped | -534.65632 | -534.44364 | -534.42971 |
| f3-4a-H-capped | -290.89432 | -290.71626 | -290.70591 |
| f3-4b-H-capped | -456.11584 | -455.95882 | -455.94769 |
| f6-7a-H-capped | -212.35615 | -212.23261 | -212.22513 |
| f7-8a-H-capped | -251.62137 | -251.47094 | -251.46197 |
| f7-8b-H-capped | -495.38605 | -495.20116 | -495.18864 |

Suppl. Table 7: **G4(MP2)-6X energies of radicals, reference structures and fragments of the deH-LNL crosslink.** The nomenclature follows Suppl. Fig. 4.

| Bond | BDE | Rad. (a) | center (a) | RSE (a) | Rad. (b) | center (b) | RSE (b) | RSE sum |
|---|---|---|---|---|---|---|---|---|
| f6-7 | 307.2 | f6-7a | N | -78.6 | f6-7b | C | -17.6 | -96.2 |
| f3-4 | 320.2 | f3-4a | C | -58.2 | f3-4b | C | -19.2 | -77.4 |
| f7-8 | 323.8 | f7-8a | C | -66.0 | f7-8b | C | -16.1 | -82.1 |

Suppl. Table 8: **BDEs, RSEs, and summed RSEs of deH-LNL bonds and radicals.** The nomenclature follows Suppl. Fig. 4. All values are given in kJ/mol. BDEs were calculated from enthalpies at 298 K (H298K).

#### 1.5 Data for peptide bonds

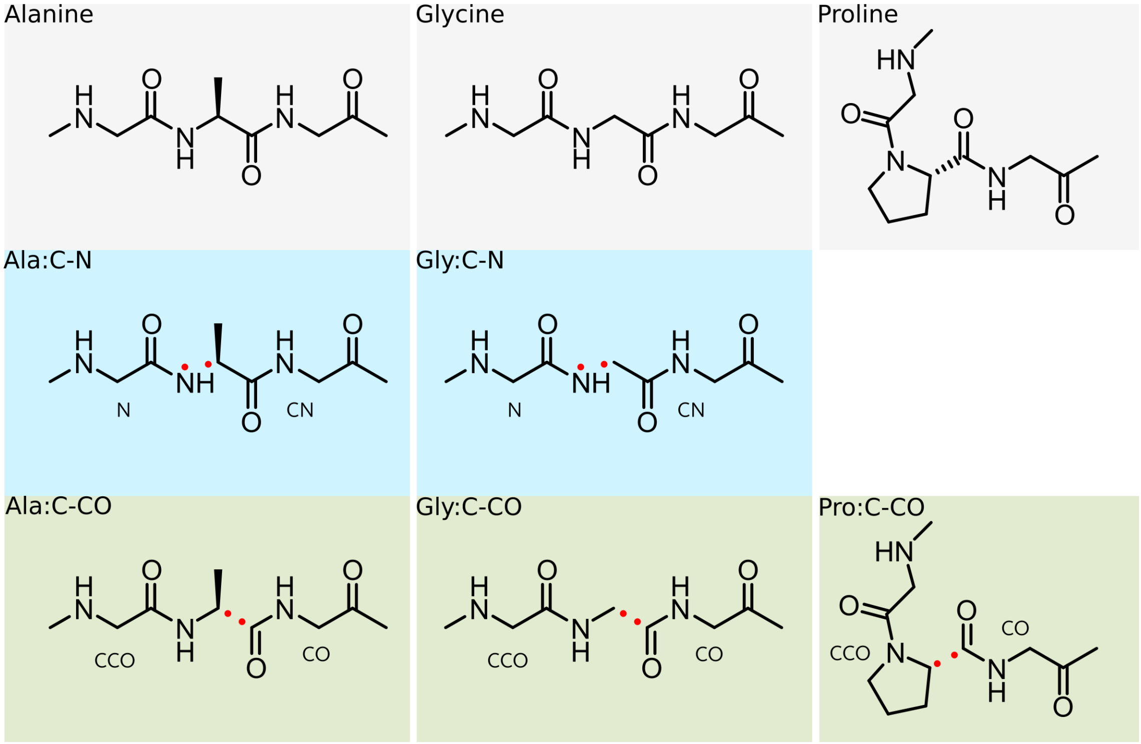


Suppl. Fig. 5: **Internal nomenclature of molecular structures used for BDE calculations of bonds in peptides.** The presented nomenclature corresponds to the Suppl. Tables 9 and 10. Bond scission of a $C_\alpha$-C bond forms the radicals CCO and CO. Bond scission of the $C_\alpha$-N bond forms the radicals CN and N.

| Molecule | ZPVE [Ha] | H0K [Ha] | H298K [Ha] |
|---|---|---|---|
| ala | -742.44664 | -742.18115 | -742.16201 |
| gly | -703.17138 | -702.93345 | -702.91566 |
| pro | -819.76896 | -819.46783 | -819.44788 |
| ala_CN | -439.39792 | -439.24643 | -439.23415 |
| gly_CN | -400.12015 | -399.99618 | -399.98560 |
| pro_CCO | -458.78656 | -458.59055 | -458.57872 |
| ala_CCO | -381.46144 | -381.30155 | -381.29019 |
| gly_N | -302.89658 | -302.79296 | -302.78475 |
| gly_CCO | -342.18689 | -342.05462 | -342.04485 |
| gly_CO | -360.84817 | -360.74991 | -360.74087 |
| ala_CN-H-capped | -440.05913 | -439.89394 | -439.88176 |
| ala_CCO-H-capped | -382.12269 | -381.94879 | -381.93768 |
| gly_CO-H-capped | -361.51025 | -361.40015 | -361.39099 |
| pro_CCO-H-capped | -459.44598 | -459.23628 | -459.22452 |
| gly_N-H-capped | -303.58727 | -303.46907 | -303.46067 |
| gly_CCO-H-capped | -342.84974 | -342.70389 | -342.69386 |
| gly_CN-H-capped | -400.78923 | -400.65175 | -400.64091 |

Suppl. Table 9: **G4(MP2)-6X energies of peptide fragments and reference structures.** The nomenclature follows Suppl. Fig. 5.

| Bond | BDE | Rad. (1) | center (1) | RSE (1) | Rad. (2) | center (2) | RSE (2) | RSE sum |
|---|---|---|---|---|---|---|---|---|
| pro_C-CO | 336.8 | pro_CCO | C | -55.0 | pro_CO | C | -43.6 | -98.6 |
| gly_C-CO | 341.2 | gly_CCO | C | -46.6 | gly_CO | C | -43.6 | -90.2 |
| ala_C-CO | 343.8 | ala_CCO | C | -50.6 | ala_CO | C | -43.6 | -94.2 |
| ala_C-N | 375.8 | ala_CN | C | -50.2 | ala_N | N | 15.5 | -34.7 |
| gly_C-N | 381.5 | gly_CN | C | -30.0 | gly_N | N | 15.5 | -14.5 |

Suppl. Table 10: **BDEs, RSEs, and summed RSEs of peptide bonds and radicals.** The nomenclature follows Suppl. Fig. 5. All values are given in kJ/mol. BDEs were calculated from enthalpies at 298 K (H298K).

#### 1.6 Details on breakage site distributions

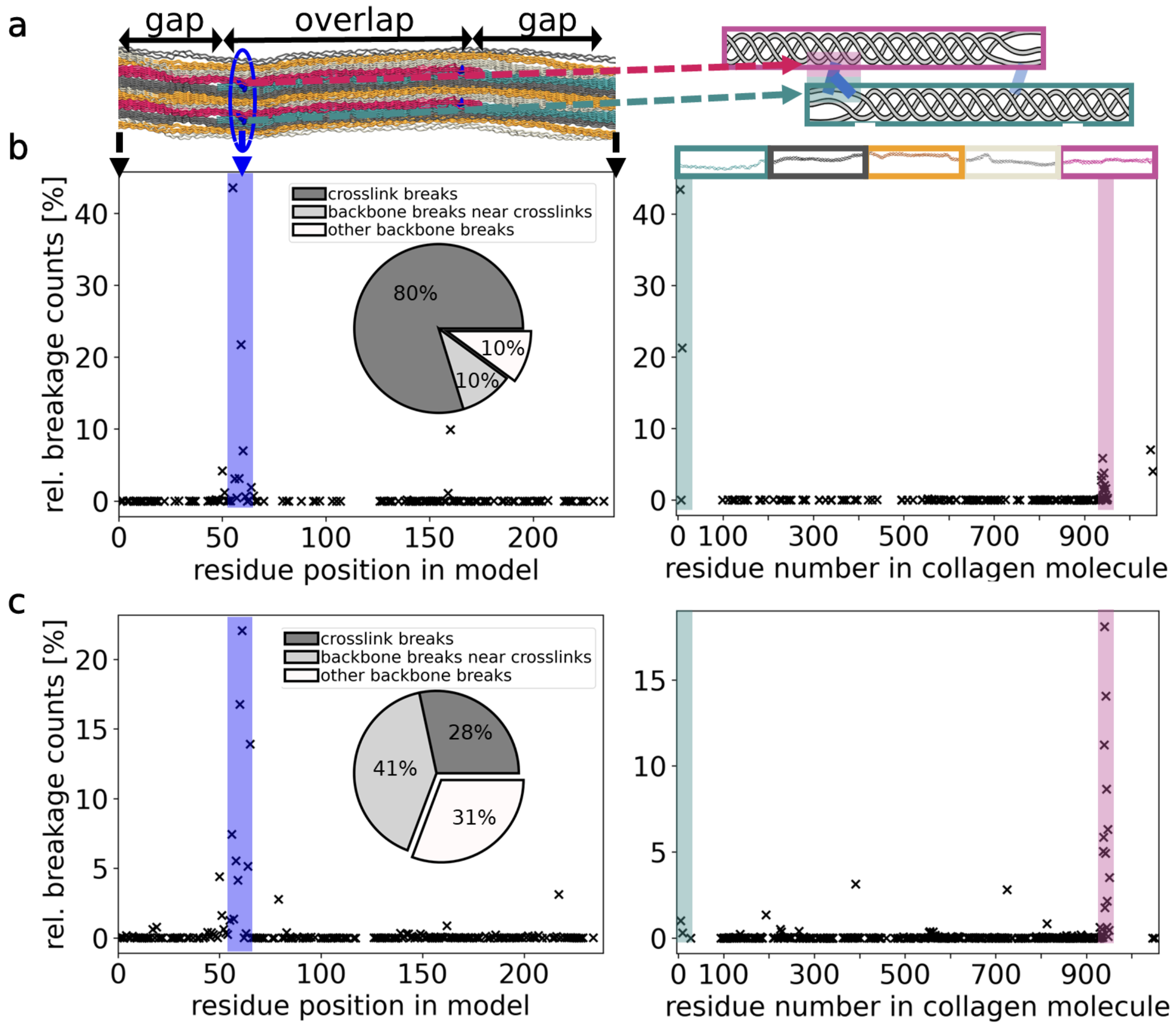


Suppl. Fig. 6: **Comparison of breakage sites in PYD using different BDEs depending on protonation states**. This analysis was performed for data of trivalent (PYD) crosslinks only. **a** Location inside the model (left) and along a collagen triple helix (right) as will be used in b and c. **b** Data for (default) BDE of 220 kJ/mol in the $C_\alpha$-$C_\beta$ bond in the shorter arm. N-terminal crosslinks as preferred breakage sites are highlighted in blue in the left panel and, in the right panel, with cyan and magenta depending on the side of the crosslink. **c** Data for BDE of 282 kJ/mol in the $C_\alpha$-$C_\beta$ bond in the shorter arm, corresponding to acidic conditions. The ruptures concentrate overall in the same regions, but are more spread out among different residues (also note the different y-axis). In the representation along the collagen molecule on the right, it can be seen how the breakages shift from one of the two arms to the leg of the trivalent crosslink.

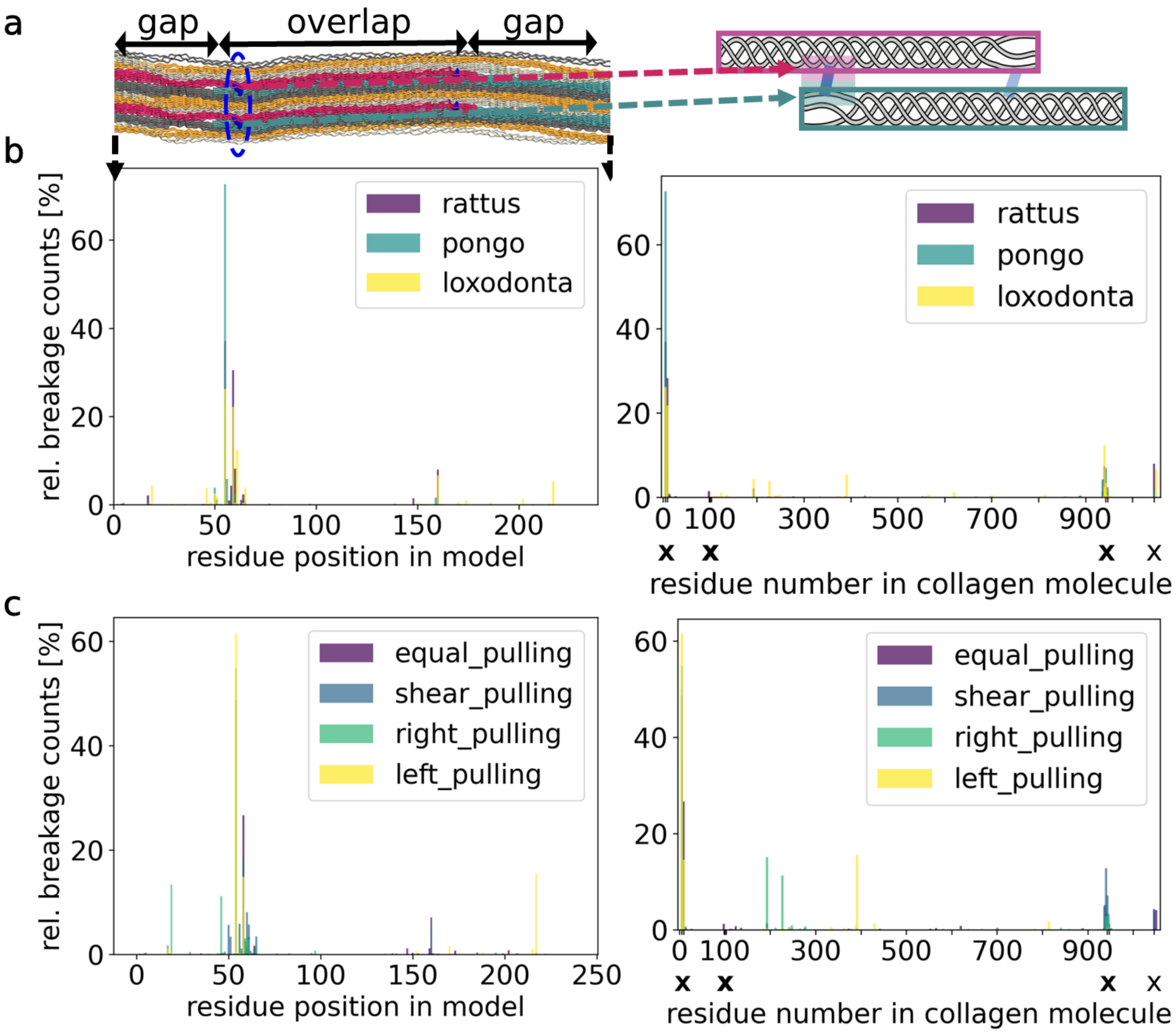


Suppl. Fig. 7: **Comparison of breakage sites depending on model species and pulling setup**. This analysis was performed for the full data set, percentages are normalized within each subset. Crosslink positions are marked with an X in the right panels. **a** Location inside the model (left) and along a collagen triple helix (right) as will be used in b and c. **b** Data for the different model species. The subsets consist of 23 simulations for Rattus, 21 for Pongo and 19 for Loxodonta. **c** Data with respect to the pulling setup. The subsets consists of 6 simulations with divalent crosslinks for each left and right pulling, 28 and 23 simulations with mixed crosslinks for equal and shear pulling, respectively.

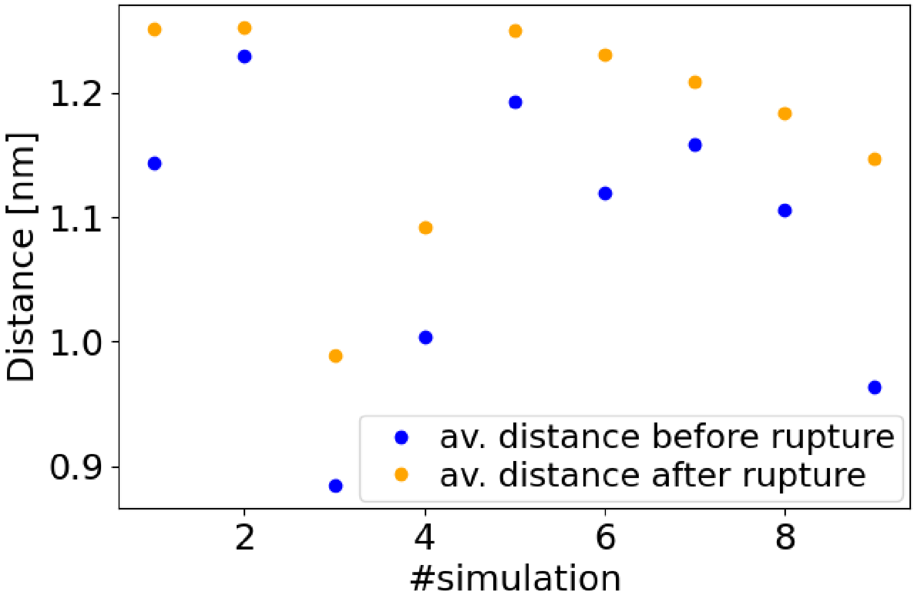


Suppl. Fig. 8: **PYD crosslinks extend after rupture of the short arm.** We measure the distance of $C_\alpha$ atoms of the (not broken) second arm to the $C_\alpha$ atoms of the leg of the PYD crosslink before and after rupturing the shorter first arm. In all replica, the distance increases indicating that the crosslink can now extend by the previously hidden length.

##### 1.7 Details on experiments

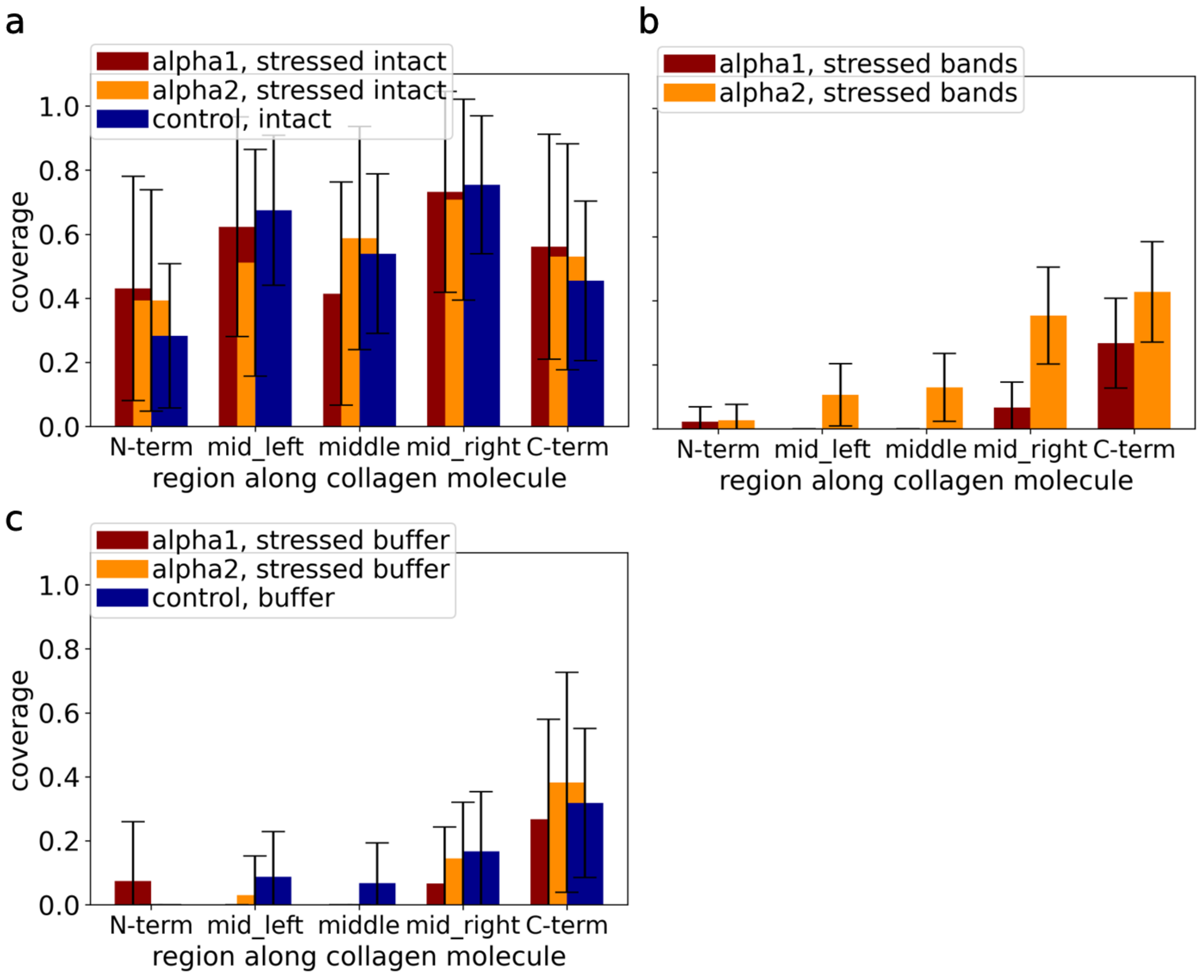


Suppl. Fig. 9: **We find collagen peptides in various areas in our SDS-PAGE gels.** Using mass spectrometry for the different bands, we find collagen fragments of both $\alpha 1$(I)- and $\alpha 2$(I)-chains from different parts along the collagen sequence. To quantify the coverage, we divided the collagen sequence in 5 regions: Before and after the N-and C-terminal crosslinks (about 100 residues each) and 3 middle parts (about 270 residues each). **a** Sequence coverage for the intact alpha-chains, both in stressed and control. **b** Sequence coverage of the 5 identified bands in the stressed samples. **c** Sequence coverage in the dye front, both control and stressed samples.

### 2 Supplementary methods

#### 2.1 Software and abbreviations

| | |
|---|---|
| **Structure design and QM visualizations** | GaussView 5.0.9 [1] |
| **QM calculations** | Gaussian 09, revision D.01 [2] |
| **MD simulations** | GROMACS, versions 2018 and 2020 [3] |
| **Bond rupture simulations** | KIMMDY [4] |
| **Visualization of molecular structures** | VMD [5] |
| **Data analysis** | Python 3.8.10, Seaborn 0.11.2 [6] |

| | |
|---|---|
| **PYD** | Pyridinoline |
| **DPD** | Deoxypyridinoline |
| **HLKNL** | Hydroxylysino-keto-norleucine |
| **deH-LNL** | Dehydro-lysino-norleucine |
| **DOPA** | Dihydroxy-phenylalanines |
| **BDE** | Bond dissociation energy |
| **RSE** | Radical stabilization energy |
| **QM** | Quantum mechanics or quantum mechanical |
| **DFT** | Density functional theory |
| **BMK** | Boese-Martin for Kinetics |
| **MD** | Molecular dynamics |
| **KIMMDY** | Hybrid Kinetic Monte Carlo/Molecular Dynamics Simulations |
| **MS** | Mass spectrometry |
| **EPR** | Electron paramagnetic resonance |

#### 2.2 Structure design and electronic structure calculations

##### 2.2.1 Design of molecular structures

Molecular structures for QM calculations were built in GuassView as shown in Suppl. Fig. 1, 2, 3, 4, and 5. By manually deleting atoms beyond broken bonds, pre-optimized radical structures were obtained from optimized molecular structures. H-capped radicals, used for radical stabilization energy calculations, were obtained by hydrogen addition to pre-optimized radical structures. In the Gaussian 09 input files, spin multiplicities were set to 1 for closed shell systems and 2 for radicals.
Naming conventions for collagen crosslinks and crosslink residues were adopted from Obarska-Kosinska et al. [7] with the exception that we extended the Greek letter notation of side-chain carbons into the aromatic system ($C_\alpha$-$C_\beta$-$C_\gamma$-$C_\sigma$-$C_\epsilon$-$C_\zeta$).
We designed the pre-optimized peptide structures such that the dihedral angles were set to common values of the respective amino acids in a Ramachandran plot of stretched fibrillar collagen type I (Rattus Norvegicus, PYD crosslink at position 9C-5B-944B (N-terminal) and 1046C-1046A-103C (C-terminal)). Initial psi and phi values were $-180°$, $-120°$ for alanine, $-180°$, $-180°$ for glycine, and $-125°$, $-90°$ for proline, respectively. The geometry of pre-optimized PYD and DPD structures was set to geometries found on the ColBuilder website [7]. In that sense, the para alkyl-substituents of the central pyridine moiety were set to be on the same side of the aromatic plane.

##### 2.2.2 Geometry optimization and single point energy calculation (G4(MP2)-6X)

We used a slightly adapted G4(MP2)-6X protocol [8] for all geometry optimizations and energy calculations. The adaptations were the following:

1. Throughout the protocol, we used finer numerical integration grids than proposed in the G4(MP2)-6X example Gaussian input shown in the supplementary file of [8]. We did so because of Bootsma et al.'s [9] findings that Gaussian 09's default numerical integration grid violates the assumption of rotational invariance.

2. The geometry optimization of the G4(MP2)-6X example input file was extended to ensure true geometry convergence and test wavefunction stability. The first geometry optimization often appeared to have converged, but more accurate force constants from the following frequency calculation showed that convergence had not yet been achieved.

3. Other than the use of finer numerical integration grids, adaptations of the energy calculation steps include calculating T1 diagnostics during CCSD(T) steps and not calculating Raman frequencies during the BMK frequency calculation.

Before energy calculation, each electronic structure was checked for convergence in all parameters and to be free of imaginary frequencies. Structures which failed to converge in their geometry were re-optimized with tighter convergence criteria and smaller optimization steps until convergeence under the initial criteria.
The final energies were calculated as described in the original paper with the provided Perl script after non-invasive modifications accounting for our input file format.
Large molecular structures (e.g., PYD substructures) had to be calculated in a two-step process due to time restraints on available computing clusters. In these cases, CCSD(T) calculations were carried out on local workstations over several days or weeks. MP2 and the following calculations were again performed on high-performance computing clusters. Consecutive log files of these calculations were prepared by concatenating both log files. Memory allocations were set dynamically.

#### 2.3 Details of energy calculations

Bond dissociation energies (BDEs) were calculated as described in Equation 1 based on G4(MP2)-6X enthalpies ($\Delta H_{298K}$) of a given molecule ($R_1$-$R_2$) and its fragments formed after homolytic bond scission ($R_1^*$, $R_2^*$). Energies of radical fragments shared by multiple structures were calculated once and reused in applicable BDE calculations. Examples are identical $C_\alpha^*$-$C_\beta$ radicals in the PYD residues LYX, LY3, and LY2.

$$\mathrm{BDE} = \Delta H_{\mathrm{298K}}(\mathrm{R}_1^*) + \Delta H_{\mathrm{298K}}(\mathrm{R}_2^*) - \Delta H_{\mathrm{298K}}(\mathrm{R}_1\text{-}\mathrm{R}_2) \tag{1}$$

In this study, radical stabilization energies (RSEs) are formally defined relative to BDEs of $CH_3$-$H$ bonds for C-centred and $NH_2$-$H$ bonds for N-centred radicals. G4(MP2)-6X enthalpies of radicals ($\Delta H_{298K}(R_1^*)$, $\Delta H_{298K}(Ref^*)$) and their H-capped counterparts ($\Delta H_{298K}(R_1\text{-}H)$, $\Delta H_{298K}(Ref\text{-}H)$) were used to obtain RSEs as given in Equation (2).

$$\begin{aligned}\mathrm{RSE}(\mathrm{R}_1^*) &= \mathrm{BDE}(\mathrm{R}_1\text{-}H) - \mathrm{BDE}(\mathrm{Ref}\text{-}H) \\ &= \Delta H_{\mathrm{298K}}(\mathrm{R}_1^*) - \Delta H_{\mathrm{298K}}(\mathrm{R}_1\text{-}H) - \Delta H_{\mathrm{298K}}(\mathrm{Ref}^*) + \Delta H_{\mathrm{298K}}(\mathrm{Ref}\text{-}H)\end{aligned} \tag{2}$$

Used reference energies were also calculated with the G4(MP2-6X) protocol:

1. $\Delta H_{298K}(CH_3^*)$ : -39.76049 Ha

2. $\Delta H_{298K}(CH_4)$ : -40.42724 Ha

3. $\Delta H_{298K}(NH_2^*)$ : -55.80836 Ha

4. $\Delta H_{298K}(NH_3)$ : -56.47839 Ha

5. $\Delta H_{298K}(H^*)$ : -0.50002 Ha

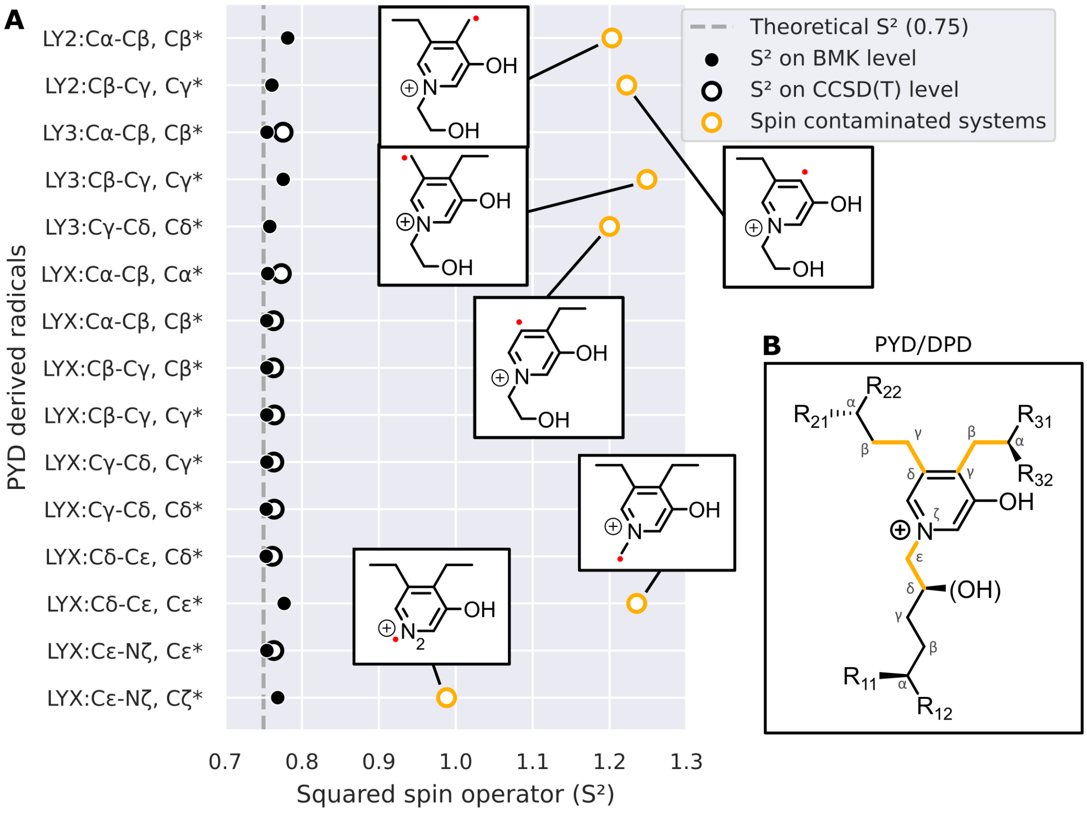


Suppl. Fig. 10: **Some electronic structures are spin contaminated in wave function-based theories.** Values of the squared spin operator ($S^2$) were automatically calculated during every G4(MP2)-6X protocol step. All shown $S^2$ values are before the annihilation of the largest spin contaminant. **A** Squared spin operator ($S^2$) for all unique PYD radicals on the BMK and CCSD(T) level. $S^2$ values on the CCSD(T) level are representative for all wave function based steps of the G4(MP2)-6X protocol. Highlighted in orange are spin contaminated structures, which all share a proximity of the radical to the aromatic system. Data points shown in black have an acceptable $S^2$ value, within the limits of $0.75 \pm 0.038$ [10]. **B** Molecular PYD and DPD structure. BDEs of highlighted bonds (orange) were calculated with a spin contaminated structure and should be carefully considered.

#### 2.4 Discussion of spin contaminated electronic structures

Suppl. Fig. 10A shows the squared spin operator ($S^2$) values of all PYD radicals, obtained on the DFT (BMK) and wave function (CCSD(T)) level of theory. In nine out of fifteen cases, both wave function and DFT $S^2$ values are within an acceptable 5 % range [10] of the theoretical value of 0.75. In six cases, however, structures became spin contaminated when transitioning from DFT to wave function theories, as done in the G4(MP2)-6X protocol. For the analysis in this manuscript, two of these six are relevant in the sense that they have BDEs low enough to compete as rupture candidates.

Spin contamination describes the artificial mixing of different spin states in calculated electronic structures, and can be an artifact of unrestricted HF-based wave functions. It can indicate that molecules require multi-reference treatments, that wave functions are unstable [10], and that electronic structures might be inaccurate. Having acceptable $S^2$ values on the BMK level, but spin contamination of the same structures on the wave function level is not uncommon for DFT functionals with low HF exchange (e.g., BMK) [11]. Interestingly, high $S^2$ values are exclusively observed for radicals in alpha or beta position to the cationic aromatic system. The radical in alpha position to the positively charged nitrogen shows lower spin contamination than other spin contaminated structures. Differing $S^2$ values between BMK and CCSD(T) wave functions indicate that on the BMK level, a doublet spin state was found as the minimal energy state, whereas on the wave function level, it was a mixture with higher-order spin states.

We tested the wave function stability of all contaminated structures on the BMK level, but could not detect any instabilities. For one electronic structure (PYD:LYX:$C_\sigma$-$C_\epsilon^*$), we further performed multiple wave function stability tests without finding instabilities. These were based on different checkpoint files of the G4(MP2)-6X protocol (BMK/6-31+G(2df,p) optimization, 2 times BMK/6-31+G(2df,p) frequency calculation, and CCSD(T)/FrzG4 energy calculation) and performed with the keyword "stable" in Gaussian on different levels of theory (BMK/6-31+G(2df,p), HF/6-31+G(2df,p), HF/6-31+G(2df,p), MP2(FrzG4)/GTMP2LargeXP, respectively). Additionally, before and after the BMK to CCSD(T) transition, the system's spin density and charge distribution were visualized but appeared to be virtually identical. T1 diagnostics [12] calculated during the CCSD(T) step were also all within acceptable limits ($< 0.045$) for open-shell systems [13]. Ideally, we would perform multi-reference calculations to validate or improve contaminated electronic structures. Unfortunately, such calculations are outside the scope of this paper. In their absence, we expect our electronic structures to be acceptable, as optimization and frequency calculations are BMK based and no wave function instabilities or suspicious spin or charge distributions were found. The fact that different spin states were found as the lowest energy states in BMK and wave function-based theories could indicate near degeneracy.

#### 2.5 Force calculation in KIMMDY: Force field influence and baseline correction

As stated in the methods section of the manuscript, we use the bond-breaking simulation scheme KIMMDY [4] and also revised the way it calculates forces during the course of this work. For this, we investigated simple polypeptide chains under force. The naive expectation here is that if pulling from the outside, the measured bond force should be the same in all amino acids inside, if we assume they behave like a chain of hookean springs. However, what we observed is a different picture: Depending on the amino acid type and position, force levels deviated measurably. Even though those differences were not large compared to their absolute levels, their error propagates exponentially in the rate (as rates are exponentially dependent on the effective energy barrier). We suspect two underlying reasons for the deviations: First, there are other bonded force field terms (angle, dihedrals) that are neglected in KIMMDY and are heterogenous among amino acids. This was corroborated by the fact that the effect varied when repeating the same calculation with Charmm36 force field [14]. This would mean that it is an artifact that should ideally be corrected for. Second, there are still other interactions ongoing in a simple polypeptide chain and the assumption of a hookean spring does not fully hold. So, what we measure are actual differences ongoing in the molecules. We also found evidence for this: Depending on the amino acids sequence in the polypeptide, i.e. the neighboring amino acids, the effects changed. This points us to the idea that the side chains might influence the also the backbone bonds, e.g. via interactions with neighbors leading to a stretch towards one direction.

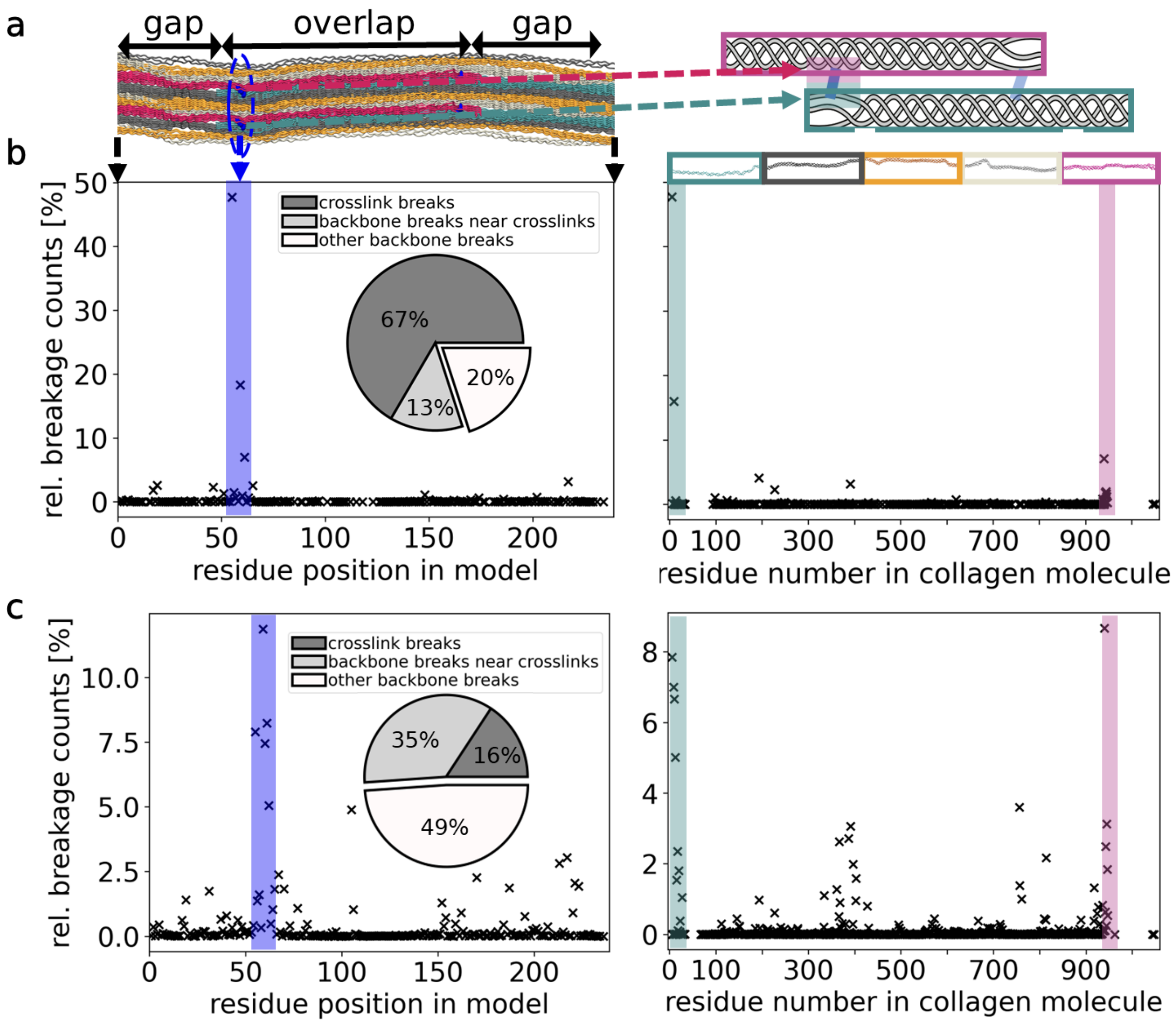


Suppl. Fig. 11: **Comparison of breakage type ratios using different methods to calculate bond forces**. This analysis was performed for data of divalent (HLKNL) crosslinks only. **a** Location inside the model (left) and along a collagen triple helix (right) as will be used in b and c. **b** Data for default force calculation. **c** Data using (maximal) correction factor. The concentration happens at the same regions, but is more spread out among different residues (also note the different y-axis).

Overall, there is probably a force-field contribution due to the nature of KIMMDY and a real contribution, which hardly can be separated by the nature of the protocol. To estimate an upper bound of the effect, we used the aforementioned polypeptide simulations as baseline reference and added a new force correction into KIMMDY, that calculates the bond forces amino acid type dependent. As this counters the effect fully, this is probably overcorrecting and mainly used for comparison in the following, which is why we provide the respective figures here in the supplement. Whereas there are quantitatively large changes on a small scale (e.g. with the correction, breakages happen in more different bonds next to each other rather in a few amino acid types where the forces used to be higher, compare Suppl. Fig. 11), already on a mid-size scale this effect seems to be small compared against the effect of the actual force distribution. So the results in this manuscript, aiming at the identification of weak regions in collagen, are robust against these considerations. Note that as stated, this is an upper limit only and the actual effect size will be smaller.
